# REV1 inhibition enhances trinucleotide repeat mutagenesis

**DOI:** 10.1101/2025.09.11.675234

**Authors:** Ava Siegel, Daniel Almstead, Naveen Kothandaraman, Jessica Reich, Erica Lamkin, Josh A. Victor, Aarzoo Grover, Kanayo Ikeh, Hannah Koval, Andrew Crompton, Hongjun Jang, Hyejin Lee, Roxana Del Rio Guerra, Dmitry M. Korzhnev, M. Kyle Hadden, Jiyong Hong, Pei Zhou, Nimrat Chatterjee

## Abstract

Trinucleotide repeat (TNR) instability has been implicated in the pathogenesis of numerous neurodegenerative disorders. Because TNR instability causes mutagenesis of the underlying gene, we refer to the repeat instability phenomenon as TNR mutagenesis in this study. While germline expansions destabilize TNR to cause disease anticipation, somatic cell TNR instability drives earlier onset of symptoms and further disease progression. However, the drivers behind these repeat length changes remain unclear. Current models suggest that DNA replication slippage events and the action of genome instability pathways, such as DNA repair, cause TNR mutagenesis. Whether mutagenic polymerases from the translesion synthesis (TLS) pathway result in TNR instability is unclear. TLS polymerases are best at bypassing difficult-to-replicate DNA regions due to bulky lesions or gaps in DNA. While some effects of TLS polymerases on TNR instability have been explored in lower organisms, evidence in human cells is lacking. Using a quantitative GFP reporter with expanded CAG repeats, we show that inhibition of the TLS polymerase REV1 by its inhibitor, JH-RE-06, or siRNA knockdown increases TNR instability and the underlying mutability. These results suggest that REV1 protects Trinucleotide repeat length mutagenesis through potential continuous DNA synthesis when replicative polymerases stall ahead of repeat secondary structures. Collectively, we present evidence of the role of the TLS pathway in TNR instability, with potential implications for understanding mutability mechanisms, disease biology, and therapeutic targeting.

## Introduction

Microsatellite repeats, consisting of tandem repeats of one to eight nucleotides, comprise around 3% of the human genome (1). The instability of microsatellite repeats is linked to over 40 diseases, most of which are neurodegenerative disorders (2). A type of microsatellite repeat, known as trinucleotide repeats (TNRs), comprises three to five nucleotide-long repetitive sequences distributed in coding and noncoding regions of the genome (2, 3). Upwards of thirteen TNR disorders are caused by the expansion of CAG repeats, including Huntington’s Disease, myotonic dystrophy type 1, and multiple spinocerebellar ataxias (4–6). These microsatellite repeats can be found at any gene region, such as the promoter, exon, intron, 5’, or 3’ area, and include trinucleotides (CAG, CTG, CGG, GAA, GCA), tetranucleotide (CCTG), pentanucleotide (ATTCT, TGGAA, TTTTA, TTTCA, and AAGGG), hexanucleotide (GGCCTG, CCCTCT, and GGGGCC), and dodecanucleotide (CCCCGCCCCGCG) (6, 7).

Trinucleotide repeat disorders follow the disease anticipation pattern, where each generation inherits expanded repeats from the previous generation. Specifically, the repeats expand in germline cells during DNA metabolic transactions. Expanded repeats cause toxic gain-of-function at the RNA or protein level (7–9). Somatic cell TNR instability has been noted in transgenic mouse models, with a tissue-specific dependence on different metabolic factors (10–14). Expanded repeats are often transcribed into unusual toxic RNA, which sequesters RNA-binding proteins or disrupts mRNA transportation, forming G-quadruplexes in nuclear foci (15–17). Other times, translation of the expanded repeat within exons causes the accumulation of abnormal poly(A) or poly(Q) tracks in respective proteins, abrogating protein function (18, 19). Curiously, expanded repeats can also initiate non-ATG translation, where repetitive polypeptides in all possible reading frames aggregate into neurotoxic clusters, contributing to disease pathogenesis (20, 21).

TNRs bear a unique architecture, where nucleotide redundancy stimulates the formation of non-B DNA secondary structures, such as intra-strand stem-loops, hairpins, H-DNA, G-4 structures, R-loops, and slipouts (6, 7, 22–24). Both germline and somatic instability of TNRs are attributed to these secondary structures (4, 25). TNR instability or mutability of the underlying gene, that is, expansion and contraction of repeat length, is attributed to mechanisms that induce DNA unwinding of expanded repeats, such as DNA replication, transcription and convergent transcription, DNA repair, recombination, and environmental stress (6, 26–33). These unwound expanded repeats tend to form secondary structures, typically stalling and restarting DNA synthesis by replicative polymerases, strand slippage or misalignment, and recruitment of other DNA metabolic pathways (6, 23, 34, 35). Specifically, the repeat secondary structures become recognition substrates for the DNA damage response (DDR) network (36–38), which is variously aided by the DNA repair pathways, such as base excision repair (BER), nucleotide excision repair (NER), and mismatch repair (MMR) causing repeat instability (6, 27, 28, 33, 39). Once engaged, each of these excision repair pathways processes the secondary structures as potential sites of damage and, by cleavage and the resynthesis steps, results in the expansion or contraction of the underlying repeats (6).

In other cases, transcriptional stalling ahead of repeat secondary structures, including R- and H-loops, causes further expansion of repeats due to break-induced replication (40). These RNA-DNA hybrids eventually produce repressive chromatin marks at and around the expanded repeat, causing local heterochromatinization via methylation and gene repression (7, 40, 41). Besides, chromatin structure and epigenetic modulation also impact the clinical severity of trinucleotide disorders via changes in gene expression and DNA methylation (25, 42–44). Bidirectional transcription (45) and a closely associated convergent transcription phenomenon are also known to contribute to TNR disease pathology via DNA toxicity and cell death mechanisms in cell cultures (31, 32).

Furthermore, in a yet-to-be-fully defined link between environmental stress and TNR instability, everyday stresses triggered TNR instability in a stress response factor and rereplication- dependent manner, where dependence on alt-non-homologous end-joining repair was observed (26, 46–48). Oxidative stress has long been shown to modulate TNR length (49); however, the connection between environment and TNR instability *in vivo* is unknown. In a yeast model, environmental stress-dependent TNR mutagenesis involved translesion synthesis (TLS) polymerases Rev1 and Pol ζ (50); the role of TLS polymerases in human cell models remains undefined. TLS polymerases are primarily known to bypass DNA damage, including playing an important role in somatic hypermutation in immunoglobulin genes and DNA repair (51–53). Importantly, TLS polymerase REV1 is recognized as the principal scaffolding protein that interacts with other polymerases to facilitate TLS activity with implications in cancer and viral biology and potential therapeutics (54–58).

Despite the range of discoveries into possible TNR instability mechanisms, therapeutic successes are minimal. Some studies focused on targeting the repeat secondary structures as therapeutic sites by using anti-sense oligonucleotides (ASOs) (59, 60), ribozymes (61), or hairpin-binding compounds, cyclic pyrrole–imidazole polyamide (CWG-cPIP) (62) to reduce the accumulation of expanded poly-Q tracks and toxic nuclear RNA foci. Other therapeutic strategies with some success are catered to symptom management, such as using pharmacotherapy (63), iron chelators (64), stem cells (65), or general toxicity reduction (66, 67). Some antioxidant therapeutics have moved into clinical trials (68–70). In this study, we tested the role of REV1 TLS polymerase in TNR-dependent mutagenesis by using a GFP-based CAG repeat assay. We found that CAG repeats were destabilized with the removal of REV1 by siRNA or by targeting it with its small-molecule inhibitor. These results establish the role of TLS polymerases in TNR instability in human cell models, with implications for understanding repeat mutagenesis mechanisms.

## Materials & Methods

### Cell Culture

T-Rex-HEK293 cells containing doxycycline-inducible GFP minigene, were grown at 37°C with 5% CO2 in DMEM with 4.5 g/L D-glucose (Gibco) supplemented with 10% (vol/vol) fetal bovine serum (FBS; Gibco) with expanded patient CAG repeats inserted within an intron of this minigene were used in this study. 0.25% trypsin-EDTA (Gibco) was used for trypsinization for culturing and cytometry experiments.

### GFP Assay

HEK293 cells containing patient (CAG)101 repeats and (CAG)0 repeats have been previously described and serve as a powerful quantitative reporter of repeat stability via changes in GFP fluorescence, where GFP fluorescence intensity tracks inversely with repeat length (71). Briefly, GFP(CAG)89 cells, which have now expanded in culture to contain (CAG)101 repeats, carry a split GFP gene under an inducible CMV/TetO2 promoter, where the cytomegalovirus immediate early promoter has two tetracycline operator sites. GFP(CAG)0 cells carry the same construct (a split GFP gene), except there are no repeats within the intron **(Supplementary** Figure 1). The expanded patient repeats embedded within the intron of the GFP gene in GFP(CAG)101 cells interfere with the intron splicing and produce a non-functional GFP gene. To quantify gene expression, cells are plated, induced with doxycycline hyclate (400 ng/ml, Sigma Aldrich), trypsinized and fixed in 1% paraformaldehyde in PBS (PFA; ChemCruz) and analyzed for GFP expression by flow cytometer (CytoFlex instrument; Beckman Coulter) with excitation through the 488 laser and emission at 525/40 filter, at the University of Vermont Flow Cytometry and Small Particles Detection (FCSPD) Facility. GFP(CAG)0 cells were used to set the gate to quantify GFP expression. Doxycycline induction produces more than 90% GFP expression in GFP(CAG)0 cells, where the GFP fluorescence peaks were separated for GFP(CAG)101 and GFP(CAG)0 cells, as shown in **Supplementary** Figure 1. Cells isolated from the right side of the gate exhibit contractions in the repeat, with approximately 7-35 repeat units remaining (72), allowing for the expression of the GFP gene and quantification of the green signal using a flow cytometer. The cells with expanded repeats in the test population are typically found to the left of the gate and can be sorted to grow clones and PCR-amplify the repeat length, as previously published (71).

GFP fluorescence is calculated as the percentage of GFP+ (GFP-positive) cells from the number of cells in the positive gate versus the total cells analyzed.

### REV1 inhibition by small molecule inhibitors

REV1 function was chemically inhibited in cells via small molecule inhibitors that target its C-terminal domain (CTD), a domain responsible for the scaffolding function for bypassing DNA damage during translesion synthesis. The scaffolding function is independent of REV1’s deoxycitidyl transferase activity, where REV1 only inserts dCTPs across DNA damages and single-strand gaps (52, 53). Mutations in the CTD’s key residues required for interaction with the other TLS polymerases abrogate TLS activity in yeast and mammalian cells (73–76); therefore, targeted drugs for this domain successfully limit the low-fidelity DNA synthesis via REV1. JH-RE- 06 (synthesized in Dr. Pei Zhou’s Laboratory, Duke University, and notated as JH in this study) targets the REV7 interface that interacts with the POLζ4 complex (the more mutagenic branch of TLS) by binding to REV1 residues and causing its dimerization that prevents Polζ4 recruitment and inhibits TLS activity **(Supplementary** Figure 2) (57). Drug 4 (synthesized by the Drs. Kyle Hadden and Dmitry Korzhnev laboratories, University of Connecticut) binds to the REV1- interacting region (RIR) region of the CTD of REV1 and limits REV1’s interaction with the RIR polymerases, POLκ, POL1, and POLη **(Supplementary** Figure 2) (77, 78). Both these drugs are specific for REV1 at low molar concentrations, and in REV1 knockout cells, there was no impact on survival or mutagenesis (57, 78).

To quantify the impact of REV1 inhibition on TNR instability, 100,000 cells were plated in 6-well plates. At 2 hours post-plating, cells were treated with 400 ng/ml of doxycycline + 1 μM JH-RE- 06 or 400 ng/ml doxycycline + 1 μM Drug 4, whereas control cells were treated with only 400 ng/ml doxycycline. At 24 hours, cells were treated with a second dose of doxycycline. At 48 hours, media was aspirated, cells were trypsinized, and fixed in PFA to be run through the flow cytometer (CytoFlex instrument; Beckman Coulter).

### REV1 inhibition by siRNA

For siRNA (small interfering RNA) knockdown of REV1, 100,000 cells were plated in six-well plates 24 hours before siRNA treatment. The next day, DMEM was aspirated out of wells, and 3 mL serum-free media was added to each control well, while 2.5 mL serum-free media was added to wells receiving the siRNA. Three sets of REV1 siRNAs **(Table 1**, purchased from Sigma Aldrich) were mixed with optimum media and incubated for 5 minutes at room temperature. Lipofectamine RNA/iMAX (Invitrogen) was mixed with optimum and incubated for 5 minutes at room temperature. The siRNA mix was added gently to the lipofectamine mix and incubated for 20 minutes at room temperature. This mix was added to appropriate wells at a final concentration of 10 nM of each siRNA. After 4 hours, the lipofectamine mix was aspirated, and 3 mL of fresh DMEM was added. This process was repeated the following day. 400 ng/ml of doxycycline was added after day 2 of transfection and again added 24 hours later, followed by GFP+ analysis by flow cytometer (CytoFlex instrument; Beckman Coulter).

**Table 1:**
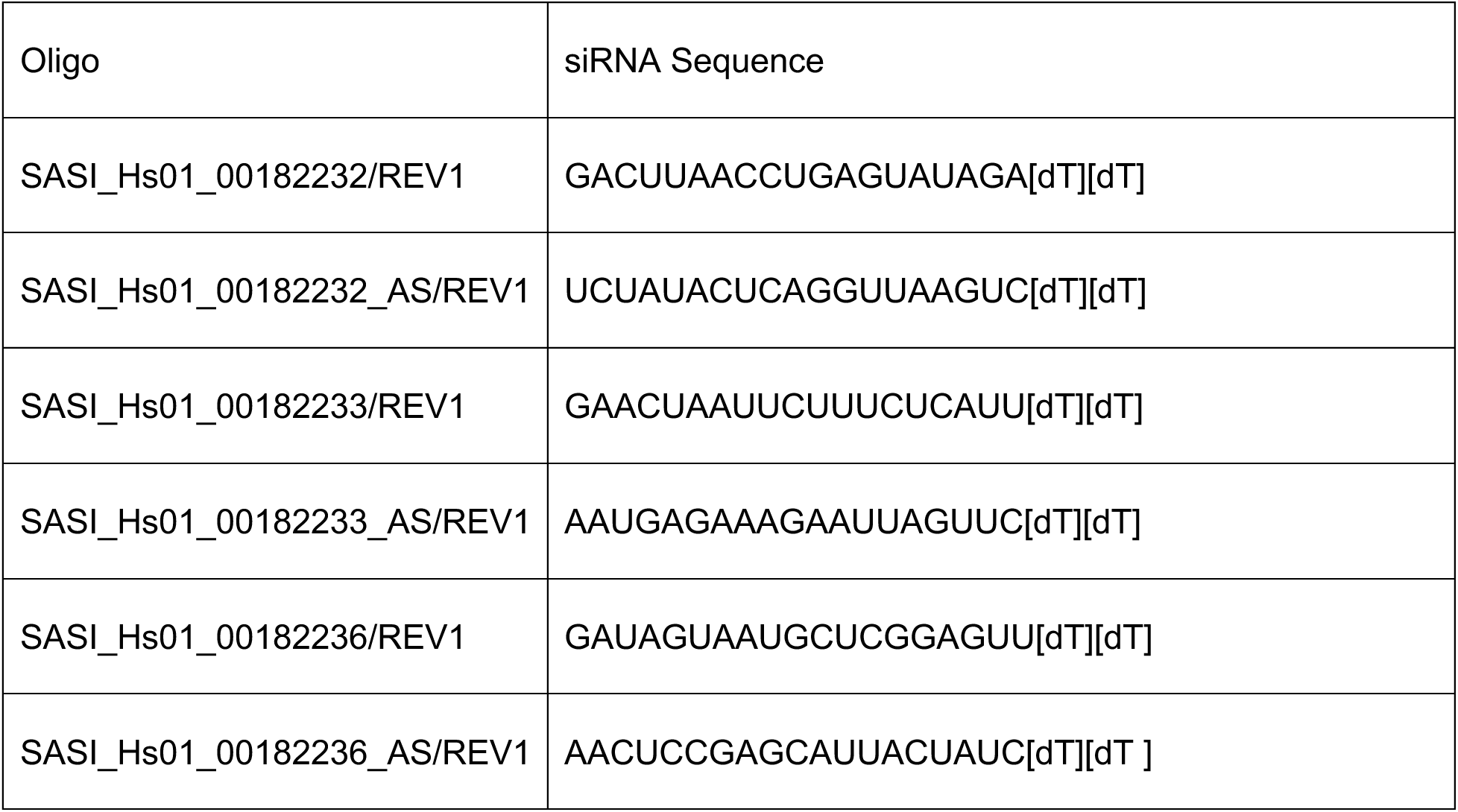
siRNA Sequences used to knockdown REV1:

### Real-Time RT-PCR

To quantify REV1 knockdown, siRNA-treated cells were pelleted, and RNA was isolated using the RNA miniprep kit from Zymo Research. RNA concentration was determined by nanodrop (ThermoFisher). About 10 ng of isolated RNA was analyzed for each reaction using the iTaq Universal SYBR Green One-Step Kit (Bio-Rad). Conditions for real-time qPCR were 50 °C for 10min, 95 °C for 1 min followed by 40 cycles of 95 °C for 15 s, and 60 °C for 1 min. The relative mRNA levels were calculated by comparing detectable products above a basal threshold, and ΔΔCT values were calculated by first normalizing values to GAPDH RNA and then to appropriate control (79). **Table 2** lists the REV1 primers used for RT-PCR reactions.

**Table 2:**
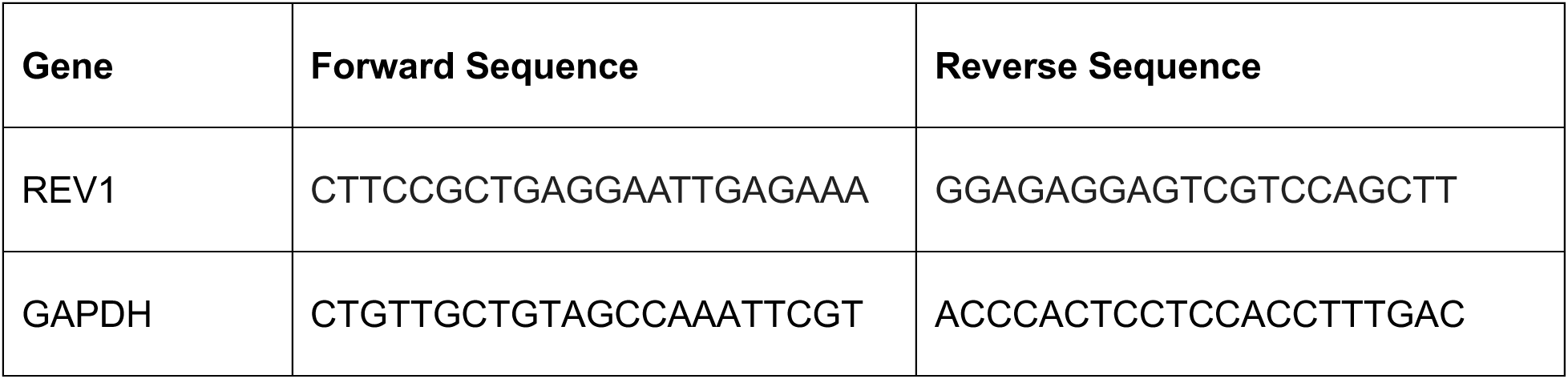
Primer sequences.

### Changes in CAG repeat length

To broadly quantify CAG repeat length changes post REV1 inhibition, 100,000 cells were plated in 6-well plates. At 2 hours post-plating, cells were treated with 400 ng/ml of doxycycline + 1 μM JH-RE-06 or 400 ng/ml doxycycline + 1 μM Drug 4 and 400 ng/ml of doxycycline alone in control cells. At 24 hours, cells were again treated with 400 ng/ml doxycycline, and at 48 hours, media was aspirated, and cells were trypsinized. DNA was isolated from a Qiagen DNA isolation kit (Qiagen). The GFP mini-gene specific primers, forward 5′- TCACAGGTTGAACCCAATCC and reverse 5′-TAAGGAGATAGTCTGCAAATTCAGTGAT, which flank the CAG repeats, were used to amplify the CAG repeat region by high-efficiency Taq polymerase (Invitrogen). PCR conditions were initial denaturation of 98 °C for 30 sec, followed by 35 cycles of denaturation, annealing, and extension at 98 °C for 10-sec min, 60 °C for 10 sec, and 72 °C for 30 sec, followed by final extension at 72 °C for 5 min. PCR products were purified using a PCR purification kit (Qiagen) and run on 1% agarose gel to confirm repeat length changes. TrackIt^TM^ 100-base pair ladder (ThermoFisher) was used to compare the band sizes.

### Statistics

FlowJo v10.10 software was used to perform the flow cytometry analysis. Student’s two-tailed t-tests were used to determine statistical significance. P values less than 0.05 were considered to be significant.

## Results

### REV1 inhibitor JH-RE-06 increases CAG repeats mutagenesis

Previously, we showed that the HEK293 cells carrying the GFP minigene with an expanded CAG patient repeat unit are an ideal model to quantify the impact of DNA metabolic processes on repeat mutagenesis (6, 26, 71). With the doxycycline-inducible, cytomegalovirus (CMV) promoter pTRE-CMV, transcription induction across the minigene reliably measures total repeat instability or mutagenesis events that are created within expanded repeats versus the no-repeat model (**Supplementary** Figure 1). Furthermore, the biological impact of mechanisms regulating the expanded repeats’ stability can be conveniently measured by functionally quantifying GFP expression through a flow cytometer.

To test the effect of the translesion synthesis (TLS) pathway, specifically REV1, which is the principal scaffolding molecule to orchestrate the low-fidelity DNA synthesis, we inhibited REV1 with either JH-RE-06 (JH) or Drug 4 in cells containing a quantitative reporter of repeat instability **(Supplementary** Figure 1). These drugs target REV1 at specific sites of its C-terminal domain (CTD) and are specific for REV1, as reported before (57, 78). JH-RE-06 binds to the REV7 interface of the CTD, causing REV1 dimerization that prevents Polζ interaction and suppression of TLS. The REV1-POLζ axis of TLS is known to cause most of the mutations. Drug 4, on the other hand, binds to the RIR interface and disrupts the interactions between REV1 CTD and the RIR-containing polymerases. The RIR polymerases can engage in redundant functions where these polymerases can successfully compensate for the absence or inhibition of any RIR polymerase, leading to less mutagenic DNA synthesis (80, 81). REV1-RIR axis is therefore considered less mutagenic.

In order to quantify the impact of REV1 inhibitors on CAG repeat mutagenesis, we exposed (CAG)101 and (CAG)0 cells to JH-RE-06 and Drug 4 for 24 hours per the experimental outline in **Figure 1A**. After a recovery of 24 hours from drug treatment and 48 hours of doxycycline induction, we fixed the cells and tested for GFP expression in a flow cytometer. We found that JH-RE-06, which targets the REV7 interface of REV1 CTD, significantly increases GFP expression compared to non-treated but doxycycline-induced (CAG)101 cells (**Figure 1B**). Here, (CAG)101 cells treated with JH-RE-06 significantly increased GFP expression. This result indicates the absence of functional REV1 and TLS activity to increase CAG/TNR mutagenesis. Surprisingly, Drug 4, which targets the RIR interface of REV1 CTD, did not significantly enhance GFP expression, suggesting that REV1-REV7, instead of the REV1-RIR axis, may be required for TNR mutagenesis. (CAG)0 cells containing zero repeats in the GFP intron showed no difference in GFP expression from REV1 inhibition. These (CAG)0 cells were used to set the GFP+ gate since the GFP exons are expected to completely reconstitute themselves into a functional protein without interference from repetitive structures (**Figure 1**). Our results in human cells confirm the findings by Collins et al. (2007), who also observed a REV1-dependent reduction in rates of repeat expansions and contractions in *S. cerevisiae* (82), suggesting a protective effect on repeat stability. In this study, JH appeared to result in a more significant increase in GFP expression, suggesting functional REV1 in human cells to stabilize TNR repeats like in *S. cerevisiae.* This study did not quantify GFP-expressing (CAG)101 cells with CAG expansions in the GFP assay.

**Figure 1:**
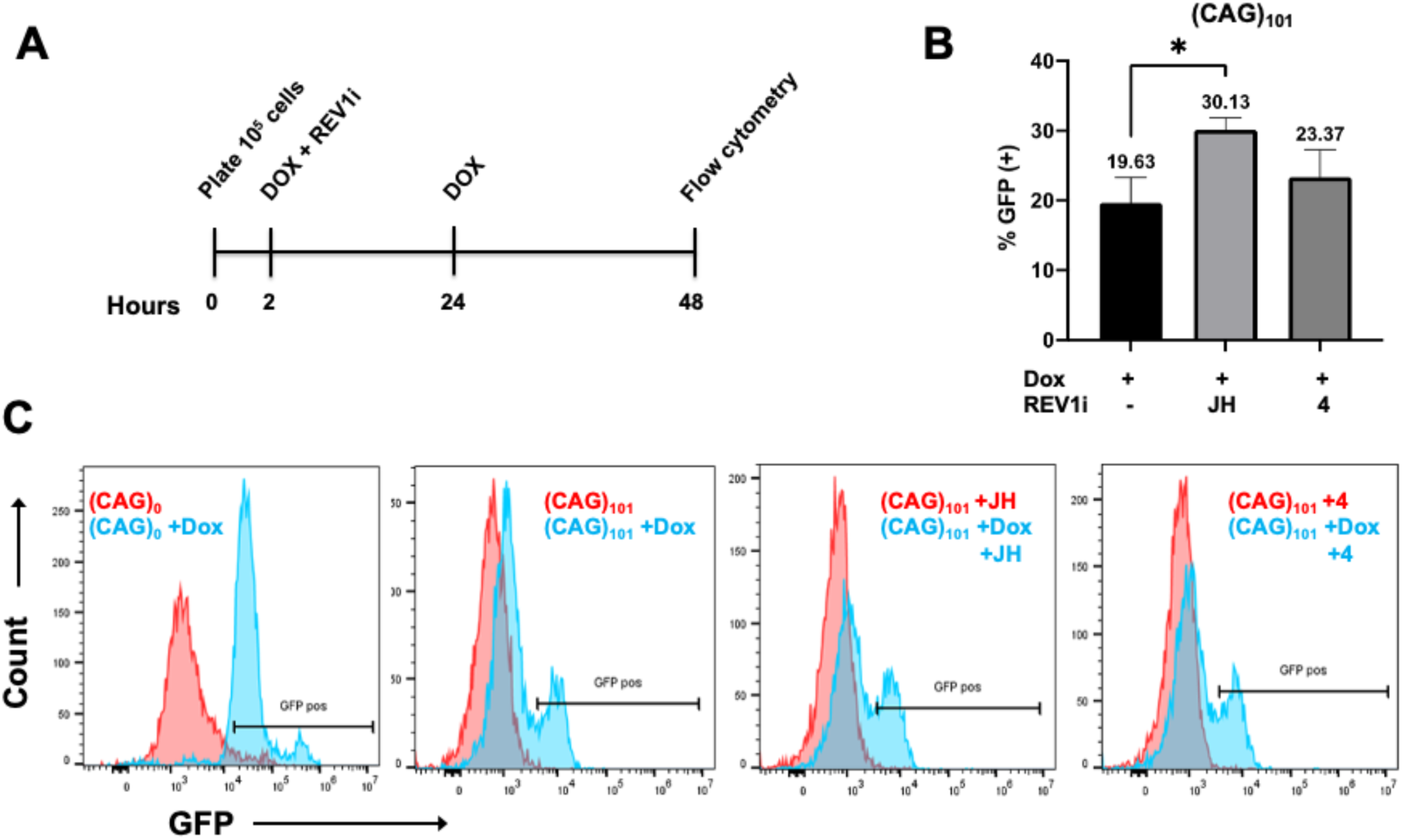
REV1 inhibition via JH-RE-06 increases TNR mutagenesis. **(A)** Experimental timeline of REV1 inhibition via REV1 inhibitors (REV1i) JH-RE-06 (JH) or Drug 4. 10^5^ HEK293 ((CAG)101 and (CAG)0) were plated in triplicates in 6-well dishes. In two hours, doxycycline at a 400 ng/ml concentration and 1 μM of JH-RE-06 or Drug 4 were added to appropriate wells. After 24 hours, a second round of doxycycline was added to the plates, followed by trypsinizing the cells, fixing them in 1% paraformaldehyde (PFA), and flow cytometry analysis at the 48-hour mark. **(B)** Percent GFP expression in (CAG)101 cells in response to indicated doxycycline and REV1 inhibitors. Percent GFP was calculated by dividing GFP-positive cells by the total number of cells analyzed in the flow cytometer. N=6, *P< 0.05, Unpaired t-test. **(C)** Histograms from flow cytometry and FloJo analysis representing GFP expression count in (CAG)0 and (CAG)101, representing data in **(B)**.

### siRNA-mediated REV1 knockdown increases CAG repeats mutagenesis

Because the REV1 small molecule inhibitor, JH-RE-06, targets only the CTD domain and induces dimerization of the protein, we reasoned that a complete knockdown of the protein would confirm the REV1 stabilizing effect on CAG repeats in (CAG)101 cells. To knockdown REV1, we used a pair of three siRNAs (**Table 1**), complexed optimum media, and the lipofectamine buffers and transfected these twice onto cells, 24 hours apart (see methods for details). Two rounds of siRNA transfection suppress gene expression better than one round. Doxycycline was added to the second round of siRNA treatment, and cells were analyzed for GFP expression using a flow cytometer (**Figure 2A and 2D**). We found that siRNA knockdown of REV1 results in more robust GFP expression than non-transfected controls (**Figure 2B**). This result supports our observation with the small molecule inhibitor of REV1, JH-RE-06, in **Figure 1B**, strongly suggesting that REV1 has a protective role in stabilizing repeats and its global removal by siRNA or inhibition by its potent inhibitor, destabilizing or mutating the repeats. Furthermore, we quantified the relative inhibition of REV1 expression in the siRNA-treated cells vs. non-treated cells. We found REV1 expression reduced by 60% via a qPCR (**Figure 2C**), confirming that the absence of REV1 increases CAG/TNR mutagenesis. As seen in the JH and Drug 4 experiments, there was also no significant increase in GFP expression of CAG0 cells after REV1 inhibition.

**Figure 2.**
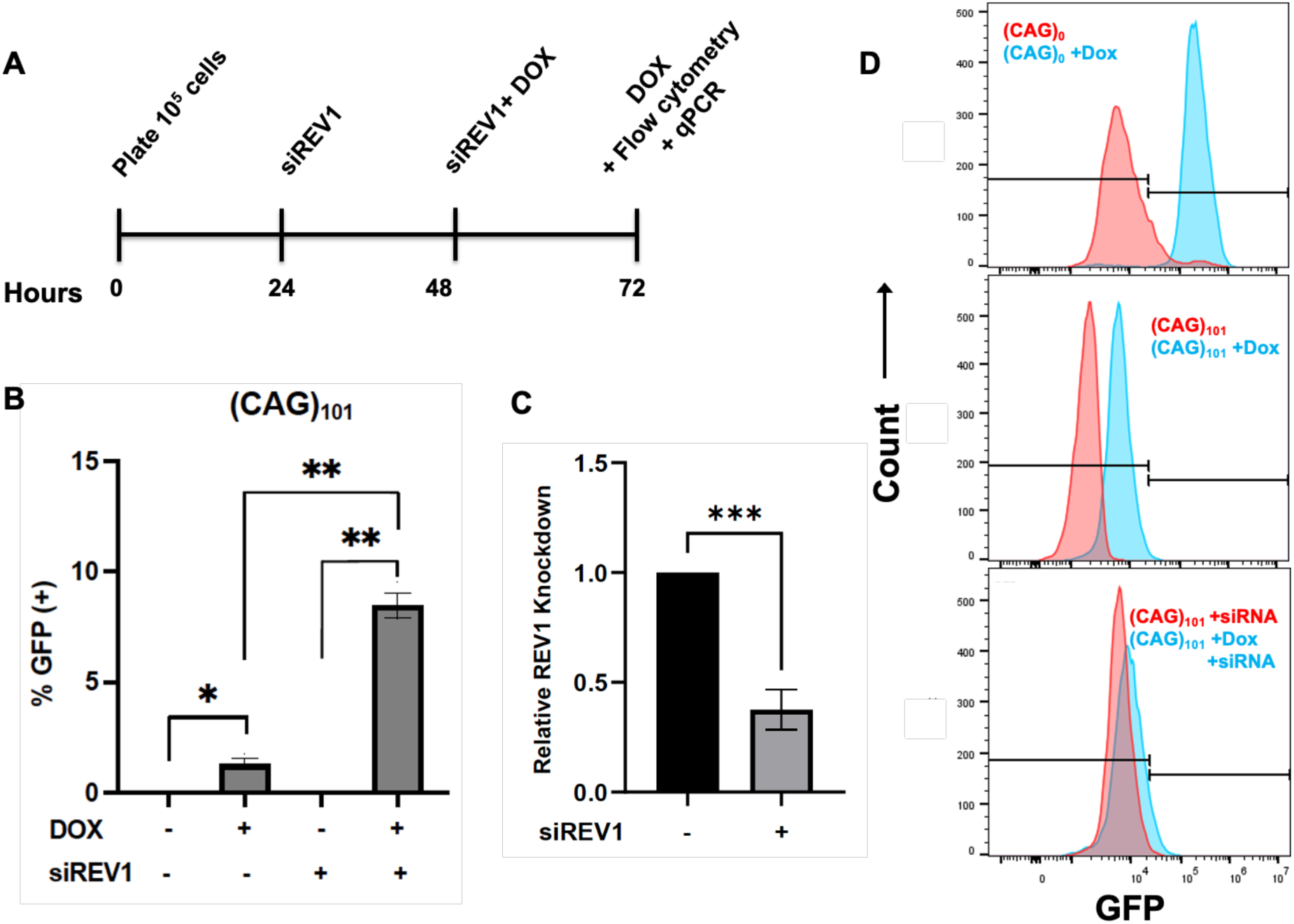
siRNA knockdown of REV1 increases TNR mutagenesis. **(A)** Experimental timeline of siRNA-dependent REV1 inhibition. HEK293 ((CAG)101 and (CAG)0) cells were plated in triplicate in 6-well dishes and, in 24 hours, transfected with siREV1. At 48 hours, a second round of siRNAs is transfected with doxycycline induction. At 72 hours, doxycycline is added again, and cells are retrieved for flow cytometry and qPCR analysis. **(B)** The representative graph shows the percentage of GFP expression in cells in response to doxycycline induction with and without siREV1 knockdown as indicated. N=2 with three technical replicates. *P< 0.05, **P< 0.01, Unpaired t-test. **(C)** The graph shows the relative expression of REV1 post-transfection with siRNA specific for REV1 as quantified by qPCR. See details in methods. **(D)** Histograms from flow cytometry and FloJo analysis representing GFP expression count in (CAG)0 and (CAG)101, representing data in **(B)**. N=3, ***P< 0.001, Unpaired t-test.

### REV1 inhibition by JH-RE-06 triggers the expansion and contraction of the CAG repeats

Since REV1 inhibition by JH-RE-06 and its siRNA-mediated knockdown resulted in increased GFP expression, indicative of increased CAG instability, we wanted to confirm this TNR instability at the level of the DNA. To do this, we treated cells with REV1 inhibitors and doxycycline, as indicated in the timeline in **Figure 3A**. At the end of the experiment, DNA was extracted and amplified by PCR per conditions detailed in the methods section. Amplification products were mixed with loading dye and run through agarose gels. **Figure 3B** shows bands specific to (CAG)0 with no repeats at around 400 bp and those within the non-treated (NT) and treated (CAG)101 cells at around 900 bp on the agarose gel. These bands set the baseline of expected sizes for the rest of the treatment conditions. Upon treatment with the REV1 inhibitor, JH-RE-06, with and without doxycycline (**Figure 3C**), we found that extra banding is indicated in JH-RE-06 and doxycycline lanes (see the lower boxed area in **Figure 3C**). Here, repeat instability is marked by the larger-sized (indicative of expansion) and smaller-sized (indicative of contraction) bands (represented in the boxes) than the 900 bp, which are absent in the control samples. These additional bands are consistent with abundant instability of CAG101 cells without functional REV1. It is to be noted that the larger-sized bands (indicated by the upper bands) in the JH-RE-06-treated samples in **Figure 3C** also appear in the doxycycline-alone-treated cells (**Figure 3B**). Still, the intensity is higher in the JH-RE-06-treated cells, indicative of more repeat instability due to REV1 limitation via this drug. Interestingly, the Drug 4 treatment did not produce the same pattern (**Figure 3D**), consistent with our results in **Figure 1B**. These results indicate that REV1 inhibition destabilizes the expanded CAG repeats, causing expansion and contraction of the resulting repeat unit.

**Figure 3:**
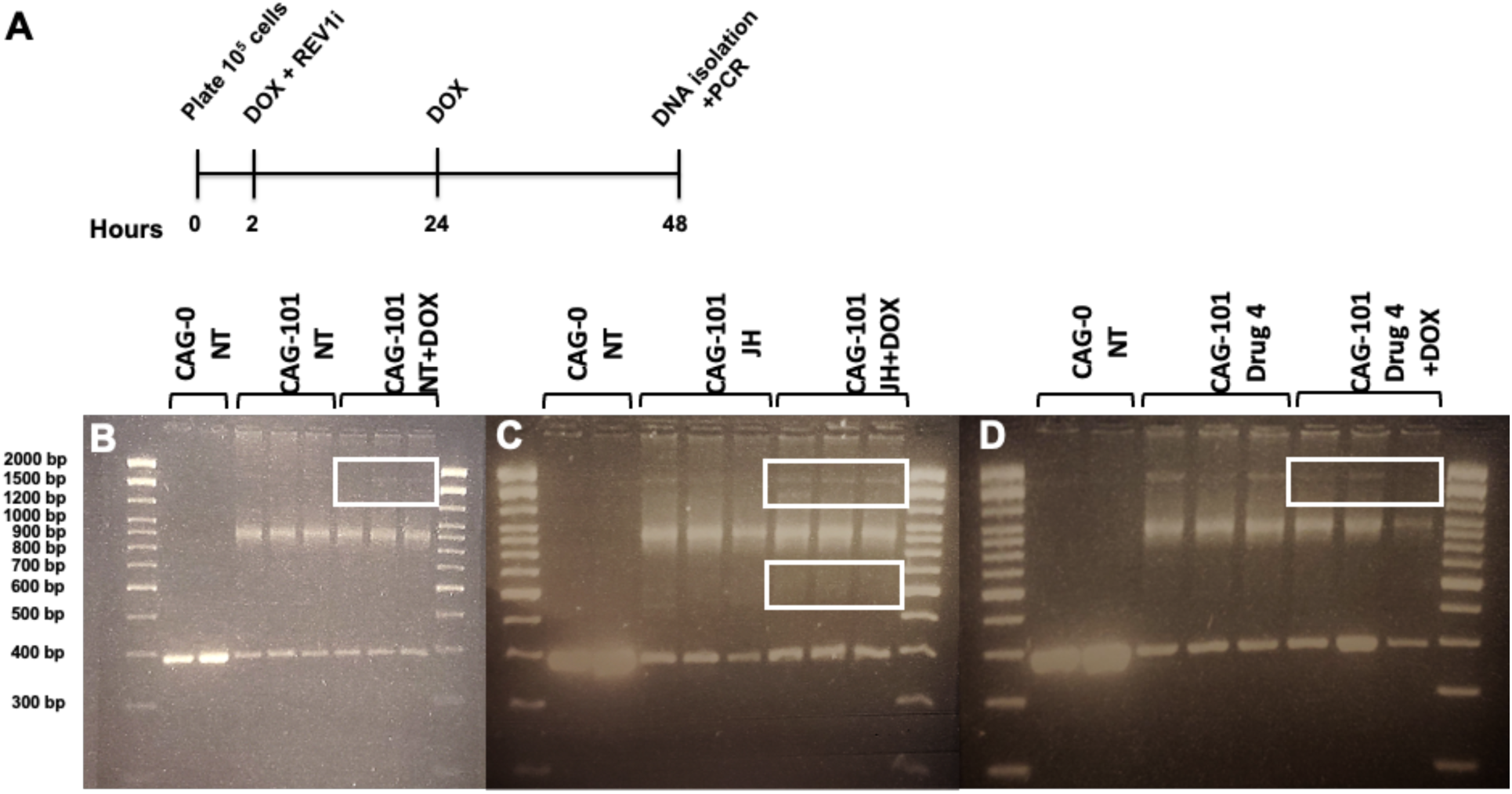
Quantification of CAG repeat length instability. **(A)** Experimental timeline to amplify CAG repeat mutagenesis by PCR. 10^5^ cells, plated in triplicate in 6-well dishes, were induced with doxycycline at a 400 ng/ml concentration, and 1 μM of JH or Drug 4 was added to the appropriate wells after two hours. A second round of doxycycline was added to the plates after 24 hours, followed by cell trypsinization, DNA isolation, and amplification via thermal cycler as described in the Methods section at 48 hours. **(B-D)** DNA gels show amplification bands (CAG)0 and (CAG)101 cells non-treated (NT) with REV1 inhibitors and 400 ng/ml doxycycline **(B)**; treated with 1 μM JH-RE-06 and 400 ng/ml doxycycline + 1 μM JH-RE-06 **(C)**; and treated with 1 μM Drug 4 and 400 ng/ml doxycycline + 1 μM Drug 4 **(D)**. Extra banding is seen in lanes of (CAG)101 cells with REV1 inhibition **(C and D)**, which is absent in non-treated cells **(B)**. TrackIt 100-base pair ladder lines the edges in all gels.

## Discussion

Translesion synthesis (TLS) is an evolutionarily conserved DNA damage bypass mechanism that utilizes specialized DNA polymerases to aid in damage tolerance by replicating over DNA lesions, thus minimizing DNA damage (52, 83). This pathway helps maintain genomic DNA integrity by rescuing stalled normal replication machinery due to DNA damage. The DNA polymerases involved in TLS belong to the Y-family (POLι, POLκ, POLη, and REV1) and B-family (Polζ) enzymes (52, 84, 85). These enzymes have specialized roles in bypassing DNA damage owing to their open structure, lack of exonuclease domain, and general low-fidelity DNA synthesis, allowing these polymerases to insert nucleotides across the damage without nucleotide complementarity during DNA synthesis. Some of these DNA damages are cognate, where a correct nucleotide is inserted across the lesion, whereas others could be bypassed in an error-prone manner, resulting in mutation formation. For instance, POLη is highly efficient in accurately replicating the cyclobutene pyrimidine dimer (CPD), the main characteristic of UV DNA damage, but highly error-prone on undamaged DNA, causing a significant increase in repeat events mutagenesis in cells (86–88). Conversely, POL1 is the most error-prone Y-family DNA polymerase and illustrates variable fidelity corresponding to the template (89). POLκ is the most widely conserved and accurate DNA polymerase among the Y-family polymerases on undamaged DNA, bypassing multiple DNA damages except for the dinucleotide lesions (90). Likewise, REV1 follows a unique, template-independent pathway to incorporate dC into the DNA template by pushing the template dG out of the helix and binding with the incoming dC to Arg324 residue through hydrogen bonding (91). POLζ4 enzyme complex includes core REV3 catalytic subunit, REV7 accessory subunit, and POL82 and POL83 subunits (92). REV7 plays a vital role in maintaining REV3 activity and binding POLζ4 to REV1 (93, 94).

During DNA damage bypass, TLS polymerases play a vital role in sustaining cell survival by engaging in low-fidelity DNA synthesis, but at the cost of undesirable mutagenesis (52, 95). Downregulation of TLS polymerases is linked to reduced mutation formation, decreased carcinogenesis, and increased sensitivity to chemotherapeutics (57, 96, 97), heralding these polymerases as key mediators in cancer etiology and subsequent treatment resistance. Mechanistically, by allowing bypass or tolerance of bulky or difficult-to-repair damages, TLS polymerases rescue stalled replication complexes, which otherwise trigger cell death. We, therefore, hypothesize that TLS polymerases are the key to rescuing stalled replication ahead of complex secondary structures formed by expanded repeats. The low-fidelity TLS pathway may be key in sustaining DNA synthesis at the secondary structures that perpetuate further expansion and lower instability, causing repeat mutagenesis at TNR loci. Trinucleotide repeats are inherently unstable due to their adopting intrastrand structural conformations identified as sites of damage by the DNA damage response (DDR) pathways, leading to their expansion and disease pathogenesis (6). In addition, other molecular processes tied to the DDR, including DNA repair, R-Loops, environmental stresses, rereplication, epigenetic modulators, and transcriptional conflicts, such as bidirectional or convergent transcription, facilitate TNR mutagenesis (6, 7, 26, 40, 41).

Studies in the yeast model highlight the engagement of the TLS pathway, another DDR and genome instability modulator, in TNR mutagenesis. For instance, Shor et al. show that removing yeast TLS polymerases Rev1 and Polζ reduced canavanine-induced mutagenesis in a run of mononucleotide repeats, which also tend to form secondary structures (6, 50). Similarly, in an elegant study, Northam et al. illustrated the formation of complex mutations from the hand-off by the stalled replication polymerases to Rev1 and Polζ that mediates an error-prone bypass of non-B DNA structures (98). Of particular interest is REV1’s role in successful G4 DNA replication via its catalytic role and POLζ4 in replication fork progression through difficult-to-replicate repetitive regions (99, 100). Repetitive DNA repeats are found at the telomeres, centromeres, and pericentromeres, possibly enriching the non-B DNA structures (101). In their expanded form in neurodegenerative disorders, trinucleotide repeats (TNRs) might similarly engage the TLS pathway during replication stalling due to non-B DNA secondary structures.

In this study, we used a fluorescent GFP minigene assay system with expanded CAG patient repeat units to quantify the impact of the TLS pathway, specifically REV1, on TNR mutagenesis. This quantitative reporter assay tracks precise changes in GFP fluorescence due to repeat length alterations, where contraction of the repeats to less than 37 are captured in the positive gate (**Supplementary** Figure 1 and (26, 71)), allowing robust identification of mechanisms propelling repeat length modifications. While the assay can capture repeat length expansions, which might accumulate within cells on the left side of the GFP histogram (**Supplementary** Figure 1), these were not studied in this study. Repeat contractions are considered reliable indicators of repeat mutagenesis as reported before (32). Hence, (CAG)101 contraction, which enables correct splicing of the repeat-containing intron and GFP expression quantified via flow cytometry, is regarded as an index of repeat mutability (**Supplementary** Figure 1). Furthermore, DNA amplification of the repeat region via a PCR conveniently captures all possible repeat length changes, providing additional means to confirm repeat length modifications comprising repeat mutagenesis (**Figure 3**). Together, this assay allows robust phenotypic identification of genetic factors that might stimulate repeat length modification.

We focused on REV1 TLS polymerase to assess the role of the TLS pathway in TNR mutagenesis. REV1 is the principal scaffolding molecule facilitating other TLS polymerases at the arrested replication site to facilitate error-prone replication (52). We used JH-RE-06, a potent inhibitor of REV1 that specifically targets the REV7 interface (**Supplementary** Figure 2), to suppress TLS activity, as previously described (57). This small molecule inhibitor is specific for REV1 while targeting the more mutagenic axis of TLS, with no off-target effects and minimal toxicity in primary cells (57). Likewise, we also tested the RIR-interface inhibitor of REV1, Drug 4, which is specific for the other interface of REV1 CTD (**Supplementary** Figure 2). Of the two small molecule inhibitors, we found that targeting the REV7-interface of REV1 resulted in more GFP expression and, therefore, TNR mutagenesis (**Figure 1**). Drug 4-dependent TNR instability was not significant. These results mirror the TLS interaction biology, where targeting the REV7 interface of REV1, the more mutagenic and consequential axis of TLS, produces more significant suppression of all TLS events versus the RIR interface (52), which is reflected in the GFP expression pattern seen in **Figure 1**.

Furthermore, siRNA knockdown of REV1 suppresses REV1 expression by about 60% and increases GFP expression, which reflects increased TNR mutagenesis due to REV1 suppression (**Figure 2**). These results confirm the role of the TLS pathway via REV1 in TNR mutagenesis. We propose that the REV1 TLS polymerase functions as a compensatory replicative polymerase at expanded TNR repeats via POLζ4 at stalled replication forks due to non-B DNA structures formed by TNR repeats (**Figure 4**). By functioning as compensatory polymerases to replicate TNR repeats complexed into secondary structures, REV1 maintains TNR length. We also propose that the REV1-dependent DNA synthesis at TNRs occur in concert with the POLζ4 complex and less so with the RIR-interface TLS polymerases, POL17, POL1, or POLκ.

**Figure 4:**
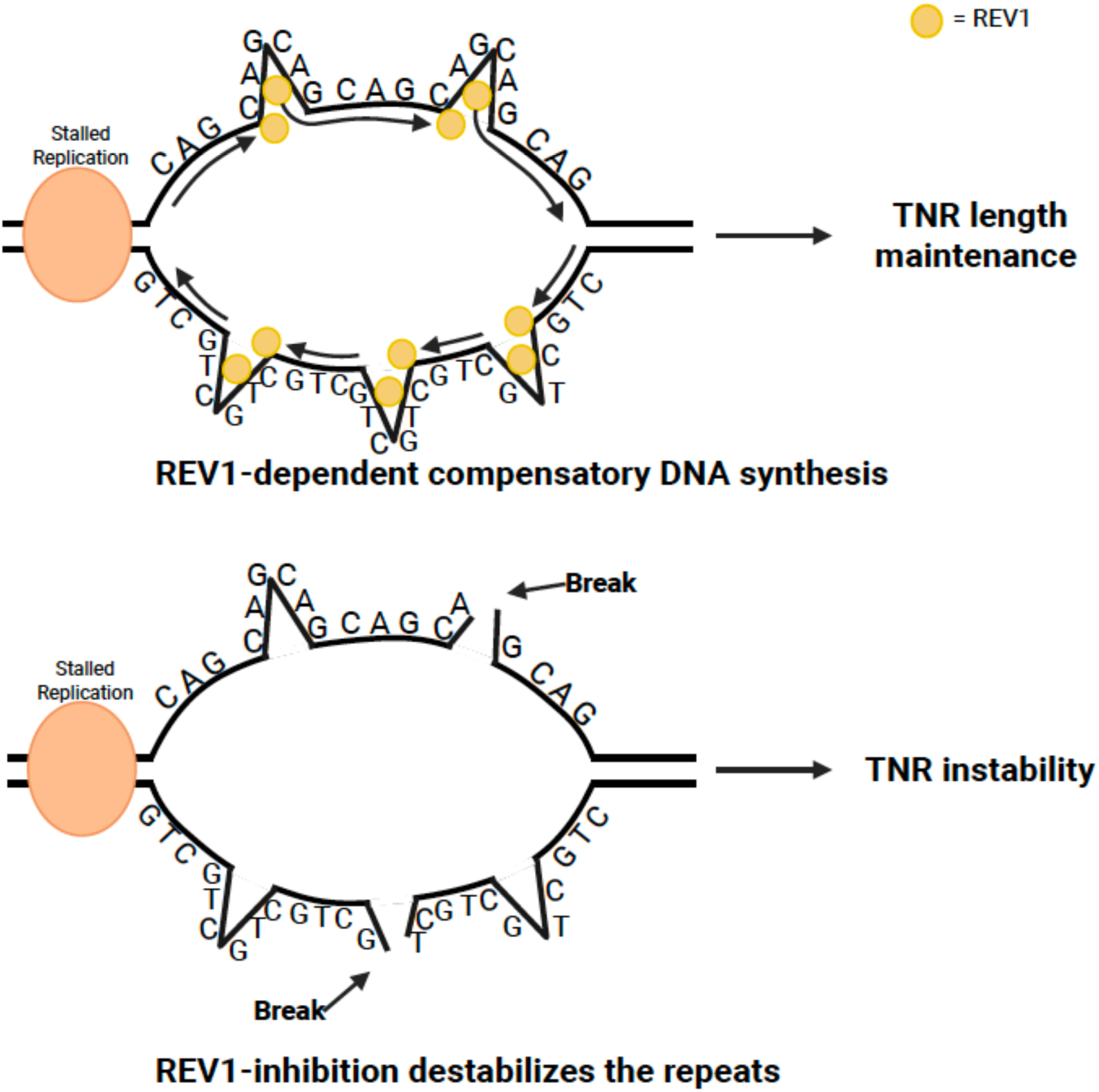
**Speculative model of REV1-dependent trinucleotide repeat stability**. During replication, the replication complex stalls ahead of unwound repetitive DNA, which engages in secondary structure formation due to intrastrand bond formation (broadly shown as triangles). REV1, due to its open structure and low fidelity, plays a compensatory role in DNA synthesis across complex repetitive DNA units, thereby limiting DNA repeat instability—expansion and contraction. In the absence of REV1, with compensatory replication absent, the replication bubble with repeat DNA-induced secondary structures becomes more destabilized and collapses, leading to repeat instability and mutagenesis.

This study shows for the first time the role of the TLS polymerases, specifically REV1, in TNR mutagenesis in human cells, with strong mechanistic implications in our understanding of neurodegenerative disorders. Because of the successful use of the REV1 inhibitor JH-RE-06 in limiting REV1 and, by extension, TLS activity in cells, future studies that utilize this drug as a chemical tool to inhibit TLS activity to study TNR disorders would be useful. Similarly, quantifying the role of the POLζ4 complex and other TLS polymerases in TNR mutagenesis will also illuminate our understanding of TLS-dependent mechanisms destabilizing TNR repeats.

This study has some limitations. For instance, although the HEK293 cell model for CAG repeat instability effectively quantifies the role of biological pathways that cause repeat length changes relevant to neurodegenerative disorders, it remains inconclusive regarding the differentiation between germline and somatic repeat mutability mechanisms. Specifically, the influence of TLS polymerases on repeat mutagenesis in germline compared to somatic cells has yet to be explored. Additionally, understanding the impact of TLS polymerases on other Trinucleotide repeats would provide insights into the mechanisms of TLS involved in non-CAG disorders. Furthermore, while the CAG repeat instability assay presented here measures repeat contraction effectively, delivering a GFP signal suitable for quantification via flow cytometry, it is limited in assessing repeat expansions, which also characterize repeat mutagenesis. Future research focused on measuring repeat expansion through fragment analysis or small-pool PCR of various repeat units, including expanded repeats, could significantly enhance our understanding of the role of TLS polymerases in TNR instability.

## Conclusion

This study presents evidence for the role of the TLS polymerase, REV1, in stabilizing TNR repeats within human cell lines. By inhibiting REV1 with small molecule inhibitors and knocking down the expression of REV1 with siRNA, we have shown an increase in TNR instability in cells lacking REV1 compared with controls. The key results in this study show a protective role of REV1 TLS polymerase in TNR stability, presenting evidence of the role of translesion synthesis and trinucleotide repeat stability.

## Contributions

A.S., D.A., N.K., E.N.L., A.C., H.K., J.A.V., J.R., A.G. and K.I., and R.D.R.G conducted cell culturing, drug treatments, PCR, DNA Agarose Gel, flow cytometry, and data analysis, V.D. HEK293 CAG cells, D.K. and K.H. Drug 4 synthesis, J.H. and P.Z. JH-RE-06 synthesis, and N.C. conceived the project, designed experiments, resources, formal analysis, supervision, funding acquisition and wrote the manuscript with help from all authors.

## Acknowledgments

We acknowledge Chatterjee lab members, especially Lindsay Allen, for their support in executing this project. This work was supported by the Larner College of Medicine Summer Research Fellowship to A.S., the Larner College of Medicine start-up funds, NIGMS MIRA R35GM150992 to N.C, and NCI R01CA233959 to D.M.K and M.K.H. We also acknowledge using the UVM-LCOM Flow Cytometry and Cell Sorting Facility (RRID: SCR_022147) support from NIH S10-ODO26843.

