## Supplementary Figures 1 and 2 for "REV1 inhibition enhances trinucleotide repeat mutagenesis"

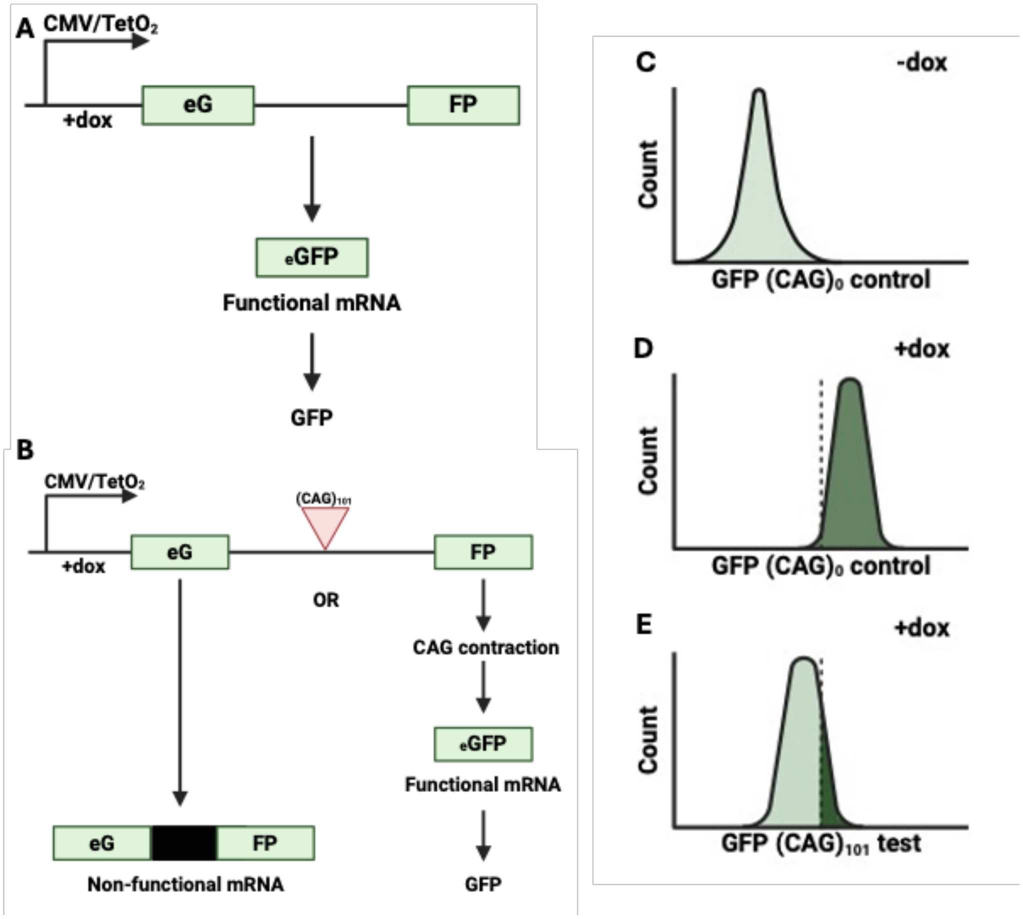


**Supplementary Figure 1: GFP Assay with flow cytometry analysis of GFP+ cells. (A)** CMV/Tet$O_2$-inducible GFP minigene construct in GFP$(CAG)_0$ cells with no repeats in the GFP intron. **(B)** CMV/Tet$O_2$-inducible GFP minigene construct in GFP$(CAG)_{101}$ cells with 101 patient-derived CAG units in the GFP intron. **(C - D)** Histogram representing the distribution of GFP$(CAG)_0$ cells in the absence and presence of doxycycline (-dox and +dox). These cellular distributions are used to set the gate. **(E)** Histogram representing the distribution of GFP$(CAG)_{101}$ cells in the presence of doxycycline. At least >94% of GFP$(CAG)_0$ cells are in the positive gate, in contrast to 1% GFP$(CAG)_{101}$ cells. Figure prepared in BioRender

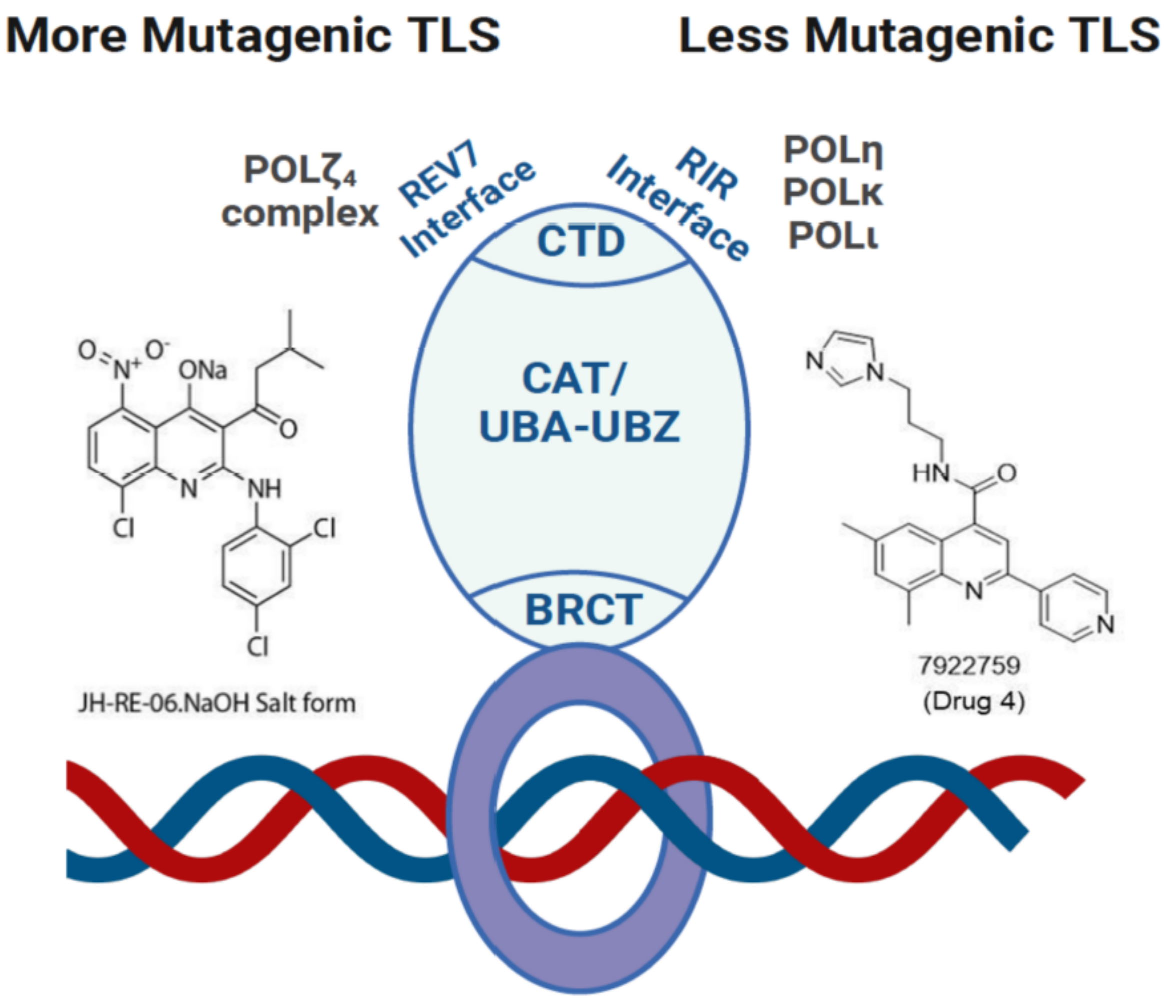


**Supplementary Figure 2: Model illustrating targeting sites of REV1 inhibitors JH-RE-06 and Drug 4.** REV1 (depicted in the blue oval), with its C-terminal domain (CTD) at the top, engages in protein-protein interactions via the REV7 interface with POL$\zeta_4$ complex and the REV1-interacting region (RIR) interface with the RIR containing translesion synthesis (TLS) polymerases, POLι, POLκ, and POLη. The REV7 interface of REV1 engages in relatively more mutability via the POL$\zeta_4$ complex, while the RIR interface results in relatively fewer mutations than the REV7 axis of REV1. Other domains of relevance to DNA damage bypass on REV1 are shown as the Catalytic domain (CAT), the Ubiquitin binding motif (UBM), and the BRCA1 C-terminal domain (BRCT) that interacts with the Proliferating cell nuclear antigen (PCNA, the purple ring) to access the DNA. JH-RE-06 (JH) binds REV1 at the REV7 interface and, by dimerizing REV1, inhibits its interaction with POL$\zeta_4$. Drug 4 binds the RIR interface of REV1 and inhibits the interaction of the RIR interface polymerases with REV1. REV1 inhibitors (REV1i)

suppress TLS activity. JH-RE-06 and Drug 4 structures derived from ChemDraw and Figure prepared in BioRender.
